# A comparison of continuous-wave fNIRS with quantitative fMRI-derived indices of brain function in the visual cortex

**DOI:** 10.64898/2026.08.10.743546

**Authors:** Giulia Rocco, Lucie Chalet, Elizabeth J. Fear, Sara Pomante, Francesca Graziano, Davide Di Censo, Manuela Carriero, Édouard Delaire, Fabrizio Esposito, Mauro Gianni Perrucci, Cosimo Del Gratta, David Perpetuini, Richard G. Wise, Antonio M. Chiarelli

## Abstract

Functional near-infrared spectroscopy (fNIRS) and functional magnetic resonance imaging (fMRI) both rely on the phenomenon of neurovascular coupling (NVC) to probe brain activity through their sensitivity to cerebral blood oxygenation. However, the relationship between fNIRS chromophores (oxy- and deoxyhaemoglobin, HbO and HbR), and fMRI (Blood Oxygen Level Dependent and Arterial Spin Labeling, BOLD and ASL) measurements, and whether this relationship remains consistent across subjects and physiological conditions, has only been partially characterised.. We acquired concurrent continuous-wave fNIRS and gradient-echo (GE) and spin-echo (SE) BOLD-ASL fMRI in healthy adults (n = 10) during visual stimulation. By applying calibrated fMRI methodology, we examined the relationships between fNIRS-derived haemoglobin modulations and fMRI-derived modulations in macrovascular (GE-) and microvascular (SE-) BOLD signals, cerebral blood flow (CBF), and oxygen metabolism (CMRO_₂_). Group-level results showed strong temporal cross-modal agreement, with HbO and HbR tightly mirroring all fMRI signal time-courses (|r| > 0.8). A quantitative analysis of trial-by-trial modulations revealed distinct state-dependent behaviours: HbO maintained a stable relationship with the fMRI-derived metrics across conditions, whereas cross-modal relationships between HbR and fMRI-derived metrics substantially strengthened at higher flow-metabolism coupling (FMC), the ratio of CBF to CMRO_₂_ change, an index of the strength of NVC. Both HbO and HbR were more strongly associated with GE-BOLD than with SE-BOLD. These findings provide a rigorous physiological grounding for fNIRS signal interpretation, demonstrating its utility as a surrogate marker for specific haemodynamic and metabolic parameters.

## 1. Introduction

Functional near-infrared spectroscopy (fNIRS) and blood-oxygen-level-dependent (BOLD) functional magnetic resonance imaging (fMRI), probe brain activity indirectly through their sensitivity to the haemodynamic response to changes in brain activity. Following an increase in neuronal electrochemical activity, accompanied by a rise in local metabolism and oxygen demand (quantified by the cerebral metabolic rate of oxygen, CMRO_2_), the phenomenon known as neurovascular coupling (NVC) increases local cerebral blood flow (CBF) and cerebral blood volume (CBV) (Roy & Sherrington, 1890; Iadecola, 2017). NVC is mediated by a complex interplay of neurons, astrocytes, pericytes, and endothelial cells, the mechanisms of which remain to be fully elucidated (Iadecola, 2017; Schaeffer & Iadecola, 2021). The ratio between the fractional change in blood flow and the fractional change in oxygen metabolism caused by varying neural activity is known as the flow-metabolism coupling (FMC) ratio. The fractional increase in CBF is generally larger than the fractional increase in CMRO_2_ leading to FMC = %ΔCBF/%ΔCMRO_2_ > 1, with values ranging typically between ∼2–3 (Fox & Raichle, 1986; Buxton & Frank, 1997; Griffeth & Buxton, 2011). This phenomenon normally results in an increase in oxyhaemoglobin (HbO) and a decrease in deoxyhaemoglobin (HbR) concentrations despite the rise in oxygen demand associated with an increase in neural activity.

fNIRS employs near-infrared light (650–1000 nm), with injection and detection points positioned on the scalp, to probe variations in brain blood oxygenation by exploiting backscattered photons that enter the cerebral cortex. The continuous-wave (CW) technology uses light sources with constant intensity and, by leveraging the differential NIR absorption of HbO and HbR, tracks haemoglobin concentration changes (ΔHbO and ΔHbR) in response to brain activity at high temporal resolution (Jöbsis, 1977; Ferrari & Quaresima, 2012). Despite being unable to quantify baseline concentrations, which would require extracting photon time-of-flight information to disentangle scattering from absorption effects (Torricelli et al., 2001), CW fNIRS is the most widely employed fNIRS technology in brain imaging, due to its simplicity, low cost, and scalability to high source-detector numerosity, covering a good portion of the scalp with a dense optode montage (Scholkmann et al., 2014). fNIRS was historically conceived as a technology probing the microvasculature, as large vessels are highly absorbing and are thought to trap photons, preventing them from reaching the detectors (Mancini et al., 1994). Beyond temporal performance and microvascular specificity, fNIRS offers practical advantages: it is portable, silent, tolerant of participant motion and free of contraindications related to metallic implants (Pinti et al., 2018; Scholkmann et al., 2014; Yücel et al., 2017), enabling deployment in settings where fMRI is impracticable (Peng & Hou, 2021). Its principal limitations are its restricted depth sensitivity (∼3 cm) (Quaresima et al., 2012), leaving subcortical structures inaccessible, and its spatial resolution (∼1 cm) which is considerably inferior to that of fMRI (Boas et al., 2004).

Blood-oxygen-level-dependent fMRI (Ogawa et al., 1990) has become the most widespread technique for non-invasive functional brain imaging. The BOLD signal arises from the paramagnetic properties of HbR, which induces magnetic field inhomogeneities and decrease the T2*-weighted MR signal (Ogawa et al., 1993). The most widely used acquisition technique is based on the echo-planar imaging (EPI) gradient-echo (GE, T2*-weighted) BOLD sequence, which is capable of producing whole-brain fMRI images with millimetre resolution and good contrast-to-noise ratio (CNR) even on clinical scanners (1.5 Tesla and higher) and has therefore broad neuroscientific applicability (Logothetis, 2008). Although GE-BOLD is the dominant approach for its simplicity and good signal-to-noise ratio (SNR), it yields a qualitative signal reflecting a composite effect of CBF, CBV, and CMRO_2_, and is highly sensitive to large draining veins, intrinsically limiting its spatial specificity (Buxton, 2009).

Complementary MRI contrasts offer additional perspectives. Spin echo (SE)-BOLD sequences employ a refocusing radiofrequency pulse that largely eliminates large-vessel contributions, isolating microvasculature signal changes at the cost of lower SNR (Uludağ et al., 2009; Yacoub et al., 2003). Arterial spin labelling (ASL) uses magnetically labelled arterial water to yield quantitative CBF estimates (Alsop et al., 2015) and can be conveniently combined with simultaneous BOLD to jointly probe cerebral physiology (Gauthier & Hoge, 2012).

Given the shared haemodynamic origins of fNIRS and BOLD signals, considerable effort has been devoted to characterising their similarities through concurrent recording studies. Early quantitative comparisons demonstrated strong correlations between GE-BOLD and optically measured ΔHbO and ΔHbR following activation (Strangman et al., 2002; Huppert et al., 2006; Steinbrink et al., 2006). Both chromophores reliably detect task-evoked haemodynamic changes, although correlation magnitude varies across studies (Scarapicchia et al., 2017; Pereira et al., 2023). Moreover, ΔHbO tracked ASL-derived CBF more faithfully than ΔHbR (Huppert et al., 2006; Hoge et al., 2005; Tak et al., 2011). However, the relationship between fNIRS chromophores and the distinct haemodynamic and metabolic parameters and vascular compartments they probe has yet to be systematically examined.

Calibrated fMRI addresses the limitations of GE-BOLD by combining BOLD and CBF measurements under isometabolic hypercapnia (induced via CO_2_ gas inhalation or volitional breath-holding, Chiarelli et al., 2007; Driver et al., 2024) to estimate the maximum possible BOLD signal change (M), which depends on baseline deoxyhaemoglobin content (Davis et al., 1998; Hoge et al., 1999). Having estimated M, fractional BOLD and CBF changes induced by a stimulus can be combined according to the Davis model to estimate fractional CMRO_2_ changes and thereby FMC ratio during changes in brain activity (Buxton et al., 2004; Gauthier & Hoge, 2012; Germuska & Wise, 2019; Chiarelli et al., 2022; Driver et al., 2024). Adding a spin-echo readout (GE–SE calibrated fMRI) provides a microvasculature-sensitive alternative to standard GE-BOLD, reducing large-vein biases and enabling a more localised assessment of cerebrovascular reactivity (Uludağ et al., 2009; Griffeth & Buxton, 2011).

To date, no study has directly compared CW-fNIRS chromophore signals to the full suite of parameters obtainable from GE–SE calibrated fMRI, leaving open the question of which cerebrovascular and metabolic parameters are most faithfully reflected by optical measures. The present study addresses this gap by acquiring simultaneous CW-fNIRS and GE–SE calibrated fMRI during a visual stimulation paradigm in healthy adults. Our primary aim was to characterise the relationships between ΔHbO and ΔHbR derived from fNIRS and the full set of haemodynamic and metabolic parameters obtained from calibrated fMRI, i.e. GE-BOLD signal, SE-BOLD signal, ASL-derived CBF, estimated CMRO_2_, and FMC, in the visual cortex. These findings are expected to provide a more rigorous physiological grounding for fNIRS signal interpretation and to inform whether fNIRS may serve as a quantitative surrogate for specific MRI-derived measures.

## 2. Material and Methods

### 2.1 Participants

Thirteen healthy subjects (two female, age = 28.2 ± 7.0) were recruited for this experiment. The study was conducted in line with the Declaration of Helsinki and was approved by the Institutional Ethics Committee of the Department of Neuroscience, Imaging and Clinical Sciences (University “G. D’Annunzio” of Chieti-Pescara, Italy). Participants provided written informed consent prior to the experiment. Data from three participants (two female) were discarded due to insufficient fNIRS data quality (see Section 2.4 for details). Therefore, data from 10 male participants were considered for statistical analysis.

### 2.2 Experimental design

Participants performed two consecutive tasks: a breath-holding task and a visual stimulation task (Figure 1a). The breath-hold (BH) task began with a 60-second baseline period, followed by 10 repeated cycles of voluntary breath-holding and a recovery period of normal breathing. The experimental timeline is detailed in Figure 1b. To increase the reliability of the BH measurement (Murphy et al., 2011), subjects were asked to perform the BH at end-expiration and to exhale completely at the end of each BH. The visual stimulation task consisted of an initial 30-second baseline, followed by 10 blocks, each comprising a task period of 31.8 s and a rest period of 30 s (see Figure 1b). During the task period, a reversing checkerboard with a fixation cross was presented (reversal frequency of 2 Hz), while only a fixation cross was presented during the rest period. The stimulation frequency was chosen as a compromise between producing a sufficient haemodynamic response in the visual cortex, of interest for the study presented here, and enabling the tracking of fast optical signals for alternative investigations (Perpetuini et al., 2026). Instructions and stimuli for both tasks were administered via E-prime (v3.0, Pittsburgh, PA, USA).

**Figure 1:**
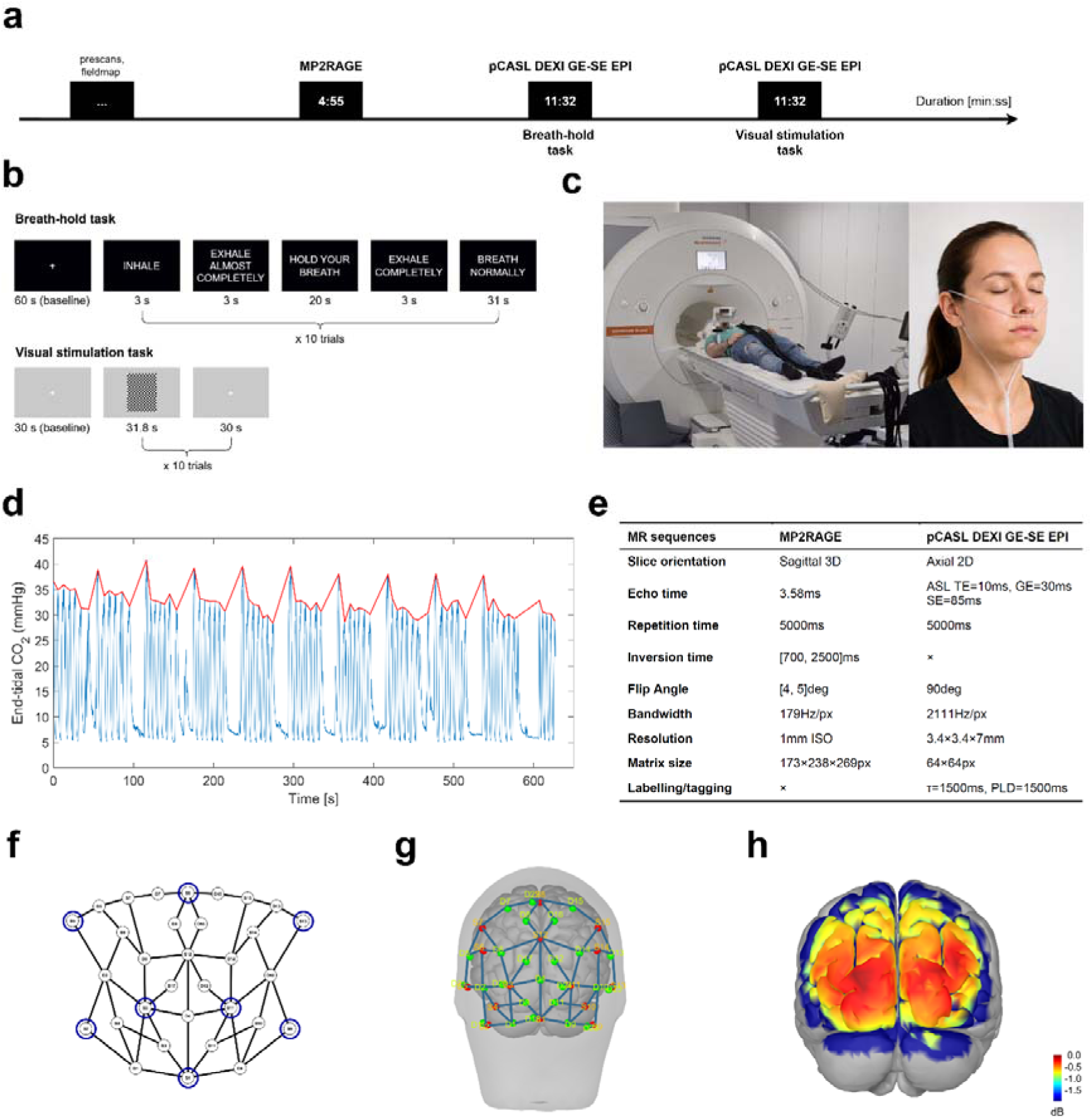
Experimental setup and data acquisition overview. (a) MRI session timeline. (b) Breath-hold and visual tasks design. (c) Simultaneous fMRI–fNIRS acquisition setup, with the participant positioned in the MRI scanner wearing the fNIRS cap (on the left) and nasal CO_2_ sampling cannula (on the right, the face is AI-generated). (d) Representative end-tidal CO_2_ (PetCO_2_) trace during the breath-hold task. Raw CO_2_ signal (blue) and the PetCO_2_ envelope obtained by interpolating expiratory peaks (red) are shown (e) Summary of MR sequence parameters. (f) fNIRS montage schematic collapsed on a 2D view. Blue circles represent short channels. (g) Optode positions projected onto a representative participant’s head surface, covering the occipital lobe bilaterally. (h) fNIRS sensitivity map (log-scale, dB).

#### CO_2_ partial pressure

During the BH task, CO_2_ partial pressure was continuously monitored by sampling expired air through a nasal cannula connected to a gas analyzer system (ML206, ADInstruments, Dunedin, New Zealand) which was in turn connected to a PowerLab 16/35 data acquisition and analysis system (PL3516, ADInstruments). Data were visualised and recorded using LabChart software (v8.1 Pro, ADInstruments). The entire experimental set-up is depicted in Figure 1c along with an example CO_2_ trace in Fig. 1d.

### 2.3 Data acquisition

#### MRI and fMRI

Data were acquired at the Institute for Advanced Biomedical Technologies, University “G. D’Annunzio” of Chieti-Pescara, Italy, on a Siemens MAGNETOM Prisma 3T scanner (Siemens Healthineers AG, Forchheim, Germany) with a 32-channel receive-only head coil.

Simultaneous BOLD and ASL data were acquired during the BH and visual tasks using an in-house pseudocontinuous ASL (pCASL) sequence based on a dual-excitation (DEXI) approach (Schmithorst et al., 2014; Chiarelli et al., 2022). The sequence employed a 2D echo-planar imaging (EPI) readout with two sequential excitations: the first with a short echo time (TE_1_ = 10 ms) optimized for ASL signal, and the second with a longer echo time (TE_2_ = 30 ms) for gradient-echo BOLD contrast. An additional refocusing pulse after the second excitation introduced spin-echo weighting at TE_3_ = 85 ms, providing sensitivity to T_2_-weighted signal changes (Pomante et al., 2026). The pCASL labelling scheme was balanced using Gaussian RF pulses (flip angle: 20°; pulse duration: 600 μs; pulse separation: 1 ms) with gradient amplitudes selected to minimize velocity sensitivity (G_avg_ = 0.8 mT/m; G_max_ = 6 mT/m; G_max_/G_avg_ = 7.5) (Zhao et al., 2017; Okell et al., 2013). Pre-labeling saturation and two inversion pulses were applied for background suppression, with the two readouts enabling optimal suppression at TE_1_ and minimal suppression at TE_2_. Both the labeling duration (τ) and post-labeling delay (PLD) were set to 1.5 s, and GRAPPA parallel imaging was used (acceleration factor = 3). The sequence timings are detailed in the Supplementary Material (Figure S1). Cerebrum coverage was achieved by acquiring 14 axial 2D EPI slices with an in-plane resolution of 3.4 × 3.4 mm² and a slice thickness of 7 mm (30% slice gap), using a TR of 5.0 s.

Two proton density (S_0_) calibration images were acquired with pCASL labeling and background suppression deactivated (TR = 7 s, TE = 10 ms), using opposite phase-encoding directions (anterior–posterior and posterior–anterior) for susceptibility distortion correction of the fMRI data and absolute ASL quantification.

A T1-weighted anatomical scan was obtained using a magnetization-prepared rapid acquisition with dual gradient echoes (MP2RAGE, Marques et al., 2010; matrix: 173 × 238 × 269, 1 mm isotropic resolution, TR/TE = 5000/3.58 ms, TI_1_/TI_2_ = 700/2500 ms) and used for image registration and brain segmentation.

Sequence parameters are summarized in Figure 1a-e.

#### fNIRS

fNIRS data were simultaneously recorded using the NIRx Borealis (NIRx Medical Technologies, Berlin, Germany), a continuous-wave fNIRS device with MR-compatible laser sources and avalanche photodiode detectors. The source wavelengths were 785 and 830 nm. The sampling frequency was 14.8 Hz. 54 long channels (with source-detector distances ranging from 21 to 42 mm) were acquired, employing in total 15 sources and 17 detectors. 8 sources were used to provide short channels (source-detector distance = 8 mm). The channel layout is shown in Figure 1f-g. In accordance with the application of visual stimulation, the montage was designed to cover the occipital lobe bilaterally. The sensitivity map is shown in Figure 1h.

### 2.4 Data processing MRI and fMRI

DICOM files were converted to BIDS by using dcm2bids (v3.2.0, Boré et al. 2023). The MP2RAGE UNI image was first denoised using MPRAGEise (v2.0, github.com/srikash/MPRAGEise), which exploits the intensity-normalized INV2 image to generate a pseudo-mask suppressing background noise, then corrected for inhomogeneity (N4BiasFieldCorrection, ANTs, Tustison et al. 2021). Tissue segmentation was performed (CAT12, Gaser et al. 2024) in MATLAB (v9.13.0, R2022b, Natick, Massachusetts: The MathWorks Inc.). The voxel-wise probability maps for each class were combined into a five-tissue label map, i.e. skin, skull, cerebrospinal fluid (CSF), grey matter (GM) and white matter (WM). GM and WM labels were merged and morphologically refined (2 mm dilation, hole-filling, 2 mm erosion) to obtain a skull-stripping mask, which was applied to the UNI image.

The skull-stripped image was coregistered to the 1-mm isotropic MNI152 atlas (Mazziotta et al. 1995) using ANTs, and the same transformation was applied to align individual T1 maps to MNI space.

fMRI data from both tasks were preprocessed identically using AFNI (Cox & Hyde 1997), FSL (Jenkinson et al. 2002), ANTs (Tustison et al. 2021), and in-house algorithms in MATLAB.

GE-BOLD series were motion-corrected (3dvolreg, AFNI), with the first volume used as the reference, and the resulting transforms applied to the simultaneously acquired ASL and SE-BOLD series (3dAllineate, AFNI). Field inhomogeneities, estimated from the S_0_ images (topup, FSL), were corrected on all three series. The GE-BOLD and SE-BOLD images were then despiked (3dDespike, AFNI). The mean BOLD image was rigidly registered to the skull-stripped MP2RAGE brain (ANTs), and the inverse transform was used to bring the CAT12-derived GM/WM maps into functional space.

The bias-field corrected S_0_ image was registered to the mean ASL time series, and was fitted with a second-order 3D polynomial to remove high spatial frequency content. Tag-control surround subtraction was then applied to the ASL volumes, which were used with the fit of the S_0_ in the kinetic model to obtain quantitative CBF (Alsop et al. 2015). The BOLD signal underwent surround averaging to eliminate perfusion contamination and was then detrended. CBF and BOLD were both expressed as relative changes from their initial baseline, estimated in the initial period at the beginning of the functional tasks. All signals were bandpass filtered (high-pass cutoff: 150 s; low-pass cutoff: 10.1 s; 4th- and 8th-order Butterworth filters respectively) and each volume was then warped to 2 mm isotropic MNI152 space with intermediate transformation to the structural T1-weighted space, which was used to compute the non-linear transformation to the template space.

Functional activation during the visual task was estimated with a subject-level GLM on SPM12 (boxcar regressor convolved with the SPM canonical HRF, six motion estimates, and a constant term), followed by a second-level random-effects GLM across subjects. Group statistical maps used one-sample t-tests, the statistical threshold was set to p < 0.05 family wise error (FWE)-corrected for multiple comparisons at cluster level with an extent threshold of 10 voxels for BOLD, while at p < 0.001 with an extent threshold of 5 voxels for the CBF. A less conservative approach was chosen for the CBF as the signal is intrinsically noisier. Clusters were labeled with a probabilistic atlas (Neuromorphometrics, embedded in SPM12).

#### PetCO_2_

End-tidal CO_2_ partial pressure (PetCO_2_), assumed to be in equilibrium with the arterial CO_2_ partial pressure, was extracted using in-house software in MATLAB (Chiarelli et al. 2022; Takano et al. 2003). The PetCO_2_ trace was obtained by identifying and linearly interpolating the expiratory peaks of the raw CO_2_ trace, as in Figure 1d. The raw PetCO_2_ signal was resampled at the fMRI TR, shifted to account for time lags between expiration and CO_2_ concentration recordings outside the scanning room, and band-pass filtered (zero-phase forward and reverse filtering with a digital IIR Butterworth filter, order 4) with cut-off times of 10.1 s (considering a Nyquist time of 10 s) and 150 s (6.7–99 mHz).

#### Calibrated fMRI framework

CMRO_2_ changes during the visual stimulation task were derived using the calibrated fMRI framework in which the breath-hold task, assuming isometabolic hypercapnia, provides the calibration step. According to the model originally proposed by Davis et al. (1998) and Hoge et al. (1999), the fractional BOLD signal change is modelled as:

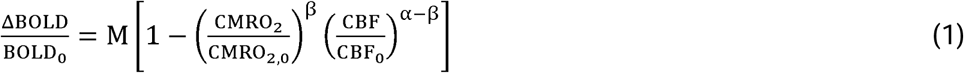

where M is the maximum BOLD signal change (in units of %) achievable at 100% venous oxygen saturation, α is the Grubb exponent relating fractional CBV to CBF changes (here set to 0.2, Chen & Pike, 2010), and β reflects field strength and vessel geometry (set to 1.3 at 3T). The subscript 0 denotes baseline values.

To estimate M, the breath-hold task was assumed to produce a purely isometabolic vasodilatory response, i.e., CMRO_2_/CMRO_2,0_ = 1 and ΔCMRO_2_/CMRO_2,0_ = 0. Under this assumption the CMRO_2_ term simplifies and M can be calculated voxelwise as:

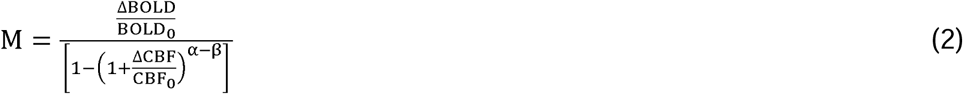

Rather than using the peak signal change, the fractional BOLD and CBF modulations induced by the breath-hold were quantified via a GLM (Chiarelli et al. 2022, Pomante et al. 2026), using the preprocessed PetCO_2_ trace as the regressor of interest for both signals (allowing a ±10 s shift, optimized by cross-correlation). The resulting GLM regression coefficients (in units of cerebrovascular reactivity, i.e., fractional signal change per unit of PetCO_2_ modulation) were then multiplied by the PetCO_2_ modulation amplitude and then substituted into Equation 2 to compute M voxel-wise.

Once the M map was estimated, it was rigidly registered to the mean BOLD image of the visual task using ANTs. Then, the CMRO_2_ change during the visual stimulation task was calculated as (Chiarelli et al., 2007):

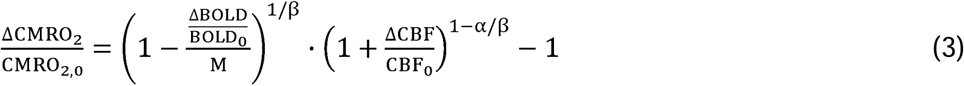

#### fNIRS

fNIRS data were processed with Homer3 (v1.33.0, Huppert et al., 2009), NIRSTORM (v0.5.1, Delaire et al., 2025) and custom scripts in MATLAB. Hereafter, we will refer to ΔHbO and ΔHbR as HbO and HbR for the sake of simplicity. Channels with a source-detector separation greater than 45 mm were discarded. Channel quality was assessed using a combination of scalp coupling index (SCI) computed as in Pollonini et al. (2016) (threshold = 0.6) and spectral power of the cross-correlated signals of the two wavelengths (threshold = 0.20). Channels with SCI or spectral power below these thresholds were pruned. For further analysis, only subjects with at least 60% of channels passing the data quality assessment were retained. Details on the subjects’ data quality are reported in the Supplementary Material (Table S2). The raw data were then converted to optical density. The motion artifacts were corrected through the Temporal Derivative Distribution Repair (TDDR) algorithm (Fishburn et al., 2019). A band-pass Butterworth filter with order 5 and cut-off frequencies 0.01 – 0.09 Hz was applied. The optical densities were then used to calculate the local oxygenation concentration changes through the modified Lambert-Beer law with subject-specific differential pathlength factors (DPF) computed according to the age-dependent formula of Scholkmann & Wolf (2013). First-level statistical analysis was performed using a GLM with ordinary least squares estimation. The design matrix included a canonical HRF regressor convolved with a boxcar function, third-degree drift regressors and the highest correlated short-separation channel for each channel. The signals were retained for further analysis after regressing out the contribution of the short-separation channels.

#### fMRI/fNIRS co-registration

Surfaces (head and cerebral cortex) were extracted during CAT12 segmentation of the MP2RAGE. To account for occasional cap placement errors where the montage extended over the cerebellum, both cerebellar lobes from the LPBA40 volume atlas (Shattuck et al., 2008) were converted into surfaces and merged with the cerebral cortex on Brainstorm (Tadel et al., 2011). We then used the scalp mesh, which captured the marks left by the sensors, since the subjects’ heads rested on the cap during scanning. Montage positions, digitized on one representative subject using a Polhemus digitizer (Polhemus Inc., Colchester, VT, USA), were mapped to each participant via anatomical fiducials (nasion, pre-auricular points) on their T1 scan, then visually fine-tuned via rigid transformation to match the sensor marks and projected onto the scalp to avoid sub-cortical coordinates.

To account for inter-subject variability in head size and cap placement, a personalised co-registration approach was adopted: only fNIRS channels located within fMRI-activated regions were retained. The group-level binarised GE-BOLD activation map (chosen over CBF to avoid overly restrictive ROIs) was warped back into each subject’s native space and projected onto individual cortical surfaces using Voronoi-based volume-to-surface interpolation (Grova et al., 2006). Montage sensitivity (Jacobian) was computed via GPU-accelerated Monte Carlo simulations with 10^8^ photons via MCXLab (Fang & Boas, 2009). The optical properties of the tissues for the MC simulations are listed in the Supplementary material (Table S1). The resulting volumetric Jacobians were interpolated onto the cortical surface using GM masking. Following exploratory data-driven simulations (Supplementary material, Section S1), we retained only the two channels per subject that exhibited the highest Jacobian sensitivity within the fMRI-activated regions.

#### fNIRS sensitivity-based amplitude correction

To account for the effect of individual anatomy on fNIRS signal amplitude (Wu et al., 2022; Yücel et al., 2025), a sensitivity-based correction was applied. Because fNIRS sensitivity to extracerebral layers varies by subject, signals were adjusted using the relationship between cortical and total tissue sensitivity. The previously calculated Jacobian was used, but three anatomical layers were considered for volume-to-surface interpolation: scalp, skull, and cortex. Surface meshes of scalp and cortex were obtained from CAT12 segmentation, while the skull mesh was generated using the SIAM model, chosen for its accuracy in segmenting skull and scalp from MP2RAGE images (Valabregue et al., 2026). For each selected channel, a correction factor was computed as the ratio of the cortical Jacobian to the total Jacobian (the sum across all three layers). The fNIRS signal amplitude was then divided by this ratio to obtain a cortically weighted signal amplitude. Further details are provided in the Supplementary Material (Section S2).

### 2.5 Statistical analysis

For each subject, fMRI signals (GE-BOLD, SE-BOLD, and CBF) were extracted voxelwise from MNI-normalized volumes, using the group-level thresholded activation map as a mask. At each timepoint, the median signal across active voxels was computed to obtain a single regional time course per subject. Fractional CMRO_2_ changes for the visual task were then estimated from these BOLD and CBF time courses using the Davis model with the median M value across the same activated voxels. fNIRS signals from the two channels with the highest Jacobian sensitivity were averaged for each subject. All signals were resampled to 200 ms time resolution and epoched into [−5, 55] s windows relative to stimulus onset (0 s).

First, we performed a temporal correlation analysis. Epoched signals were averaged within subject (median across trials) and across subjects (mean), and Pearson’s correlations were computed both within and between modalities at the group level, then repeated at the individual level using each subject’s task-evoked average (median across trials) prior to group averaging. Negative HbR (−HbR) signal was considered for this analysis.

A trial-by-trial analysis was then performed to test whether a quantitative, rather than purely temporal, relationship existed between fNIRS and fMRI signals. In other words, we investigated whether the magnitude of a given BOLD (or CBF, or CMRO_2_) fractional change in a trial corresponded to the magnitude of the concurrent fNIRS response. To mitigate inter-trial variability and noise, each subject’s z-scored averaged time series served as a regressor for that subject’s single trials of the same modality (e.g., BOLD trials were regressed against the subject’s average BOLD response). This yielded a β-weight per trial, reflecting the single-trial response magnitude relative to the subject’s mean. Correlations between β-weights were computed across all signal pairs. Given the repeated-measures data structure (10 trials × 10 subjects), the significance of each correlation was assessed using repeated measures correlation (Bakdash & Marusich, 2017). The resulting p-values were corrected for multiple comparisons using Benjamini-Hochberg false discovery rate (q < 0.05). Finally, we investigated whether correlation coefficients varied systematically with the FMC, splitting the trials into two subgroups using the global median FMC as a threshold. Negative HbR (−HbR) signal was considered for this analysis.

## 3. Results

### fMRI activation maps

Group-level activation following visual stimulation was consistently detected in the primary visual cortex across all three fMRI modalities (GE-BOLD, SE-BOLD, and CBF), as shown in Figure 2. Cluster statistics are detailed in the Supplementary Material (Table S3). The activated clusters were bilateral and located in the posterior occipital region, consistent with the expected visual areas to the stimulation task. However, distinct differences in cluster extent and statistical sensitivity were observed across the modalities. Specifically, GE-BOLD showed the largest cluster volumes across the left and right inferior occipital gyri (> 500 voxels, > 4000 mm^3^) yielding peak t-values up to t = 20.14, with additional significant activations in adjacent structures. SE-BOLD displayed a more spatially constrained activation pattern centered around the primary visual area (approximately 500 voxels). CBF maps showed a high spatial overlap with the BOLD modalities, but with characteristically smaller cluster sizes (up to 100 voxels) and lower peak statistical values (maximum t = 7.15).

**Figure 2.**
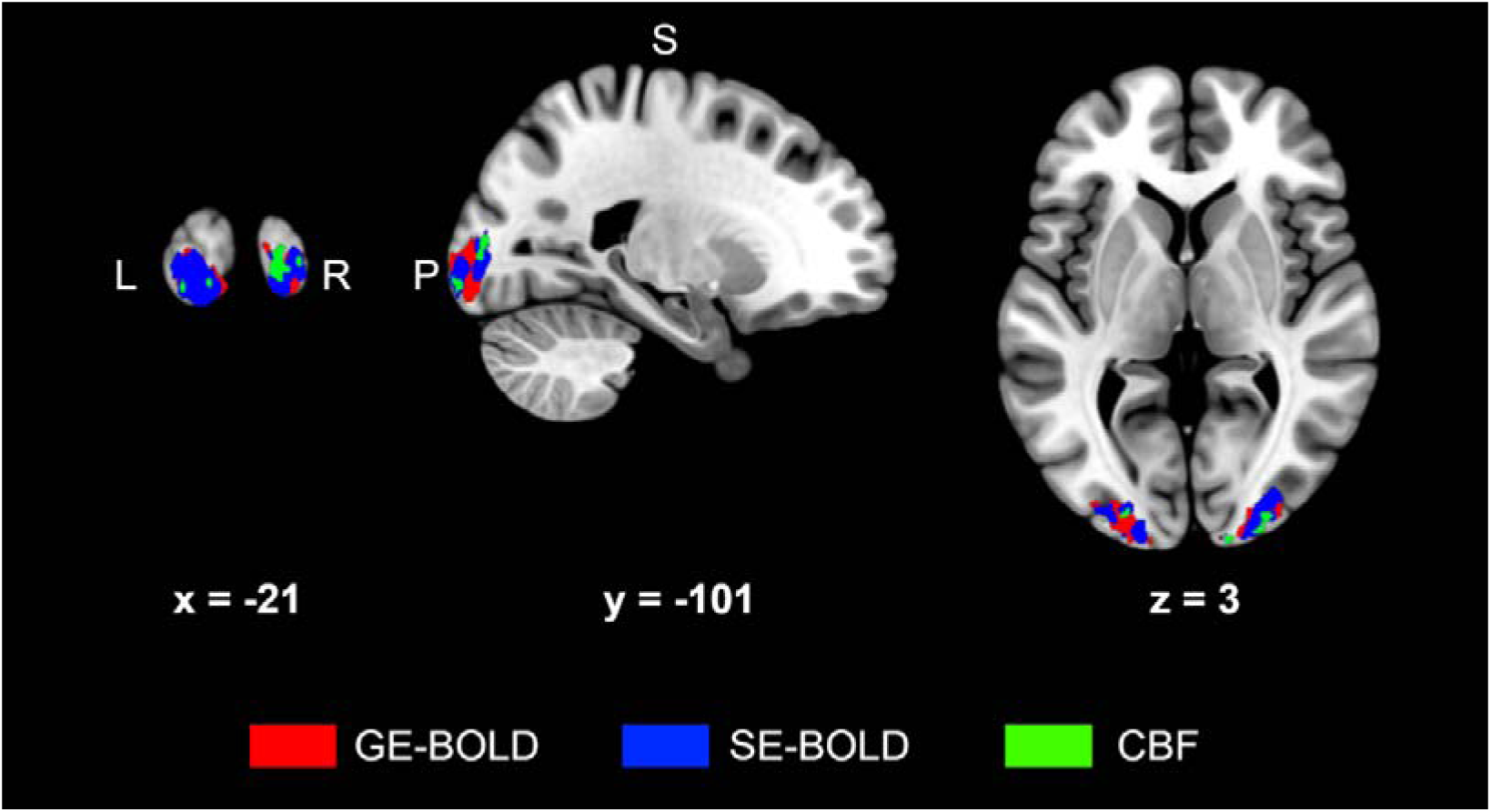
Group-level fMRI activation maps for the visual stimulation task. Thresholded statistical maps for GE-BOLD (red), SE-BOLD (blue), and CBF (green) overlaid on the MNI152 template. Thresholds: GE-BOLD and SE-BOLD at p < 0.05 FWE-corrected (cluster extent ≥ 10 voxels); CBF at p < 0.001 uncorrected (cluster extent ≥ 5 voxels). MNI coordinates are millimetres. L: left; R: right; S: superior; P: posterior.

### Temporal correlation analysis

Group-averaged task-evoked responses in the activated areas are shown in Figure 3a. All fMRI signals exhibited a clear positive haemodynamic response peaking between 15 and 20 s after stimulus onset. The fractional signal changes reached peak means of 0.95 ± 0.26 % for GE-BOLD, 0.73 ± 0.24 % for SE-BOLD, 39.03 ± 9.81 % for CBF and 22.82 ± 4.50 % for CMRO_2_. fNIRS signals showed a concurrent positive HbO response peaking at 0.91 ± 0.76 μM, a corresponding negative HbR response peaking at -0.39 ± 0.36 μM and HbT response peaking at 0.52 ± 0.47 μM.

**Figure 3.**
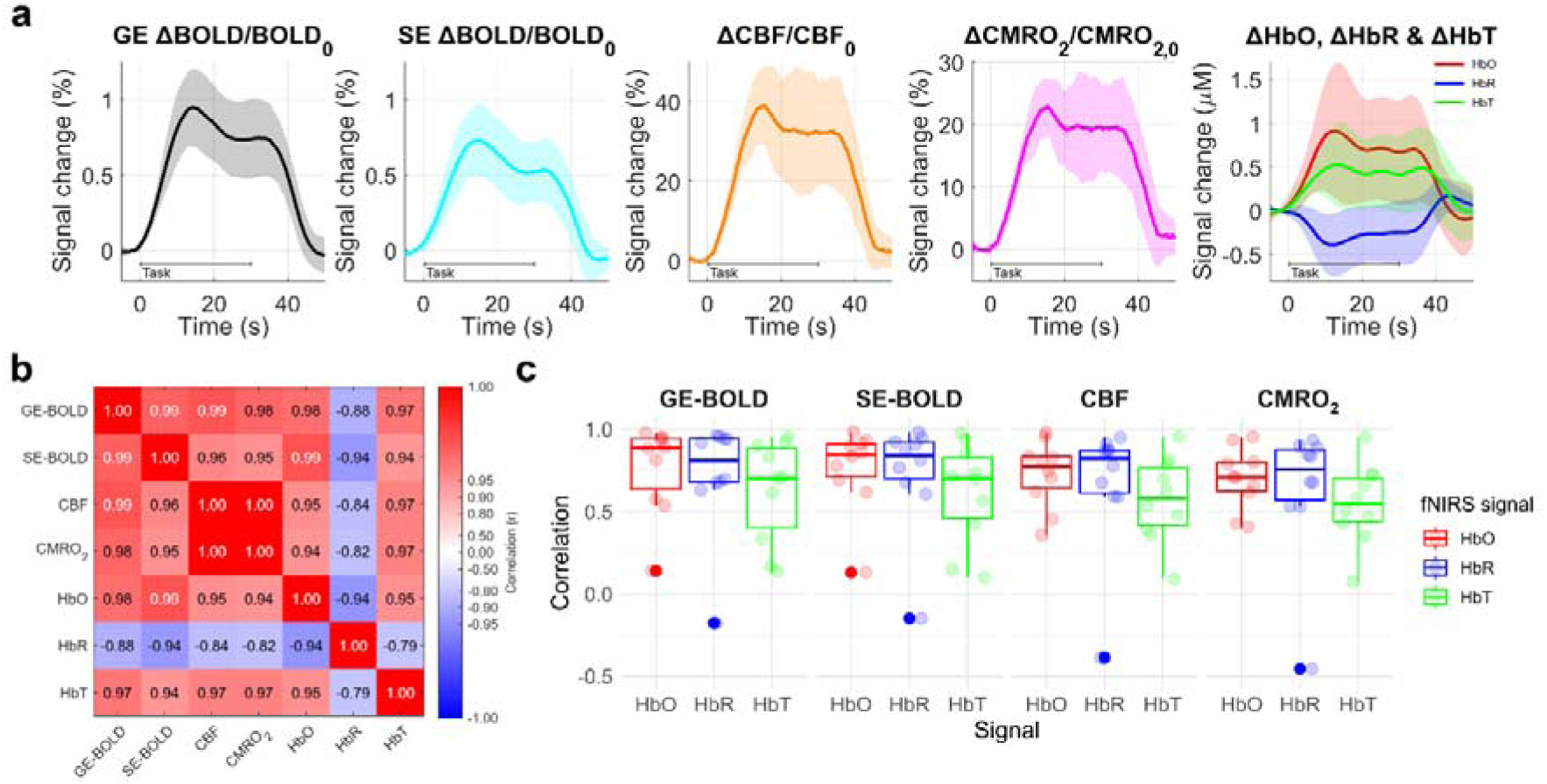
Group-level temporal responses and cross-modal temporal correlations during the visual stimulation task. (a) Group-averaged task-evoked time courses for fMRI GE-BOLD (black), SE-BOLD (cyan), CBF (orange), CMRO_2_ (magenta), fNIRS HbO (red), HbR (blue) and HbT (green). Solid lines represent the mean across subjects; shaded areas indicate ± standard deviation. (b) Group-level Pearson correlation matrix computed between the mean time courses in (a). Color indicates Pearson’s correlation coefficient (r), rendered on a sign-preserving logarithmic scale [sign(r)·−log₁₀(1−|r|)] to visually separate strongly correlated values (|r| > 0.8). Colorbar ticks are labeled with the corresponding untransformed r-values (c) Individual-subject Pearson correlation coefficients between each fNIRS chromophore (HbO, red; HbR, blue; HbT, green) and each fMRI-derived signal (GE-BOLD, SE-BOLD, CBF, CMRO_2_). Each dot represents one subject.

The group-level temporal correlation matrix (Figure 3b) revealed very high within-modality correlations across all fMRI signals (r ≥ 0.95 for GE-BOLD, SE-BOLD, CBF and CMRO_2_). Cross-modal fNIRS-fMRI correlations were also strong: HbO demonstrated strong positive correlations with all fMRI metrics (r = 0.94 to 0.98) together with HbT (r = 0.94 to 0.97), while HbR consistently exhibited strong negative correlations (r = −0.82 to −0.94).

At the individual subject level (Figure 3c), Pearson correlation coefficients between fNIRS and fMRI time courses remained predominantly high across subjects, confirming the stability of these cross-modal relationships despite expected inter-subject physiological variability. Both HbO and the inverted HbR signal showed consistently positive correlations with all fMRI modalities (median r = 0.71 to 0.89 for HbO; 0.76 to 0.84 for HbR; 0.55 to 0.71 for HbT). One subject showed outlying correlation values, likely reflecting suboptimal fNIRS signal quality, potentially affected by extracerebral confounds. Except for the outlier, all the correlations were highly significant (p < 0.001).

### Trial-by-trial analysis

The trial-by-trial distribution of modulation amplitudes, extracted as beta values for each signal metric across all trials (num trials = 100, 10 trials x 10 subjects) is summarised in Figure 4a. fMRI metrics yielded beta values of 0.36 ± 0.13 % (mean ± std) for GE-BOLD, 0.29 ± 0.12 % for SE-BOLD, 14.54 ± 7.72 % for CBF, and 8.51 ± 4. 95 % for CMRO_2_. For the fNIRS metrics, the mean concentration changes were 0.41 ± 0.36 μM for HbO, 0.21 ± 0.15 μM for HbR, and 0.28 ± 0.26 μM for HbT. FMC showed a mean value of 1.86 ± 0.98, with a median value of 1.68, which was used as a threshold in the subsequent analysis to form two subgroups with the same number of trials. Two outliers (one with FMC > 5, the other with FMC < 0) were not shown for visualization purposes. The proximity of the mean (diamond) and median (bar) values across the fMRI and fNIRS metrics in Figure 4a indicates generally symmetric trial distributions, with minor positive skewing in HbR and HbT likely attributable to lower SNR and cross-metric dependency. Furthermore, the uniform dispersion of individual trial data points (i.e. evenly spread colour points) across all response ranges confirms that the quantitative modulations reflect true trial-by-trial physiological variance and that there is not a specific subject skewing the distribution.

**Figure 4.**
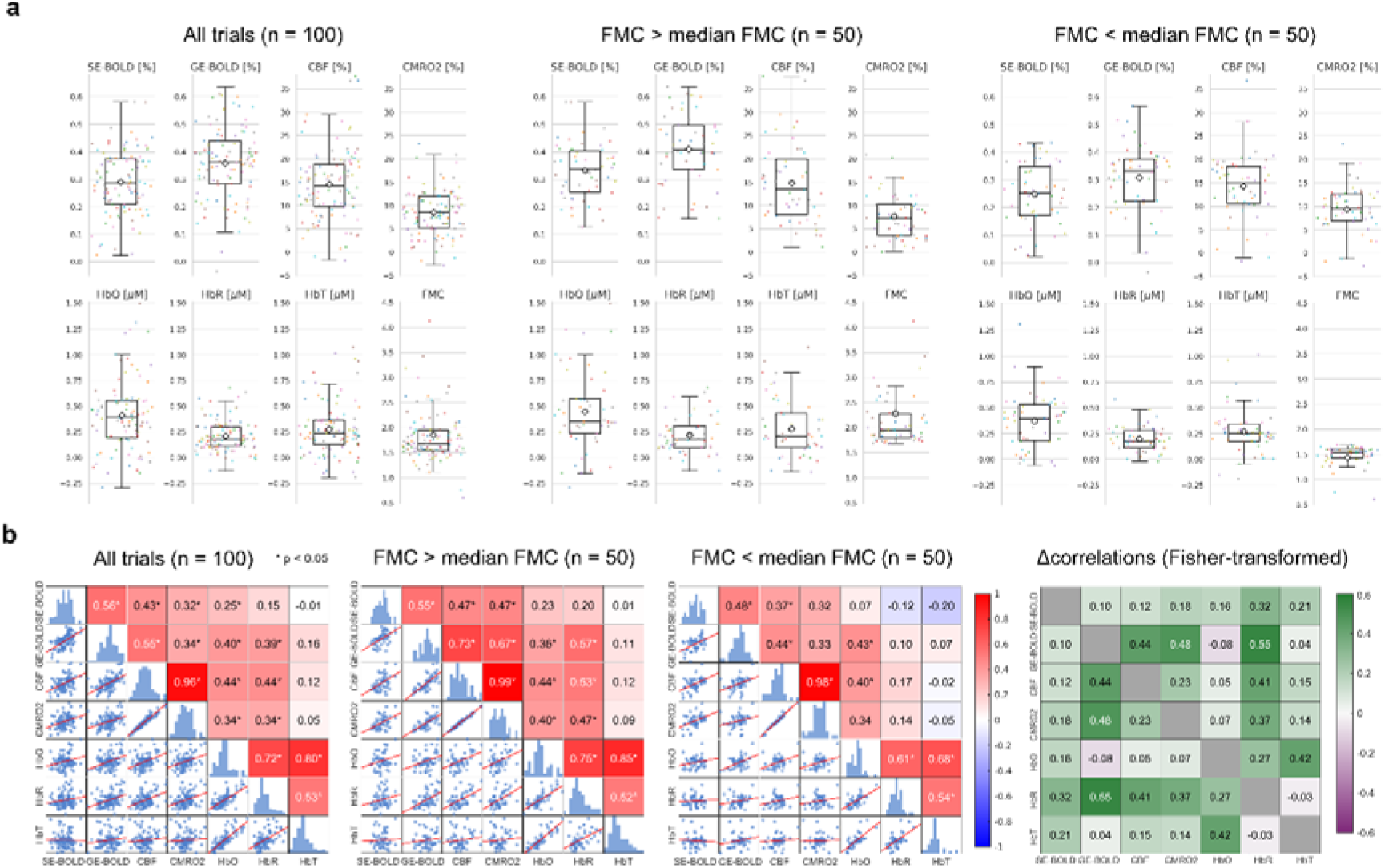
Trial-by-trial response amplitudes and cross-modal quantitative correlations. (a) Distribution of single-trial beta weights (response amplitudes) for each signal metric and FMC across all trials (n = 100, 10 trials × 10 subjects, left), high FMC trials (FMC > median, n = 50, center), and low FMC trials (FMC < median, n = 50, right). Box plots show median (bar), interquartile range, and whiskers extending to 1.5× IQR; diamonds indicate the mean; colored dots represent individual trials, with each color corresponding to one subject. (b) Pairwise Pearson correlation matrices of single-trial beta weights for all signal pairs, computed across all trials (left), high FMC trials (FMC > median, n = 50, center), and low FMC trials (FMC < median, n = 50, right). Asterisks indicate statistically significant correlations (p < 0.05). Scatter plots below the diagonal show individual trial data; histograms on the diagonal show each signal’s marginal distribution. The matrix on the right represents the difference between the Fisher-transformed correlations of the high and low FMC groups.]

In the trial-by-trial correlation analysis between the beta values relative to the quantitative modulations (Figure 4b), several statistically significant positive correlations (q < 0.05) emerged among the signals. With all trials combined (n = 100), cross-modal correlations were generally moderate. CBF and CMRO_2_ showed moderate correlations with HbO (r = 0.44 and 0.34, respectively, q < 0.001), and similarly with HbR. GE-BOLD correlated similarly with HbO (r = 0.40, q < 0.001) and HbR (r = 0.39, q < 0.001). SE-BOLD showed the weakest cross-modal coupling overall, correlating significantly only with HbO (r = 0.25, q < 0.05). None of the MRI parameters correlated significantly with HbT.

Stratifying trials into high FMC (FMC > median, num trials = 50) and low FMC (FMC < median, num trials = 50) subgroups (median FMC = 1.68) showed that FMC modulated coupling with HbR, while HbO coupling was relatively independent. Correlation between GE-BOLD and HbR increased from 0.39 (all trials, q < 0.001) to 0.57 in the high FMC subgroup and dropped to non-significant (r = 0.10, q > 0.05) in the low FMC subgroup. To better identify the FMC modulations between subgroups, the difference between the Fisher-transformed correlations of high and low FMC (Δz) was visualized as a difference matrix (Figure 4b, on the right). HbR and GE-BOLD showed the largest FMC-driven shift (Δz = 0.55). A similar pattern held for CBF–HbR (0.53 high vs. 0.17 low, Δz = 0.41) and CMRO_2_–HbR (0.47 high vs. 0.14 low, Δz = 0.37). In contrast, correlations with HbO remained relatively stable and largely significant across subgroups (CBF–HbO: 0.44 vs. 0.40; CMRO_2_–HbO: 0.40 vs. 0.34; Δz ≤ 0.07). SE-BOLD and HbT showed weak, mostly non-significant coupling with the other modality regardless of FMC state.

The correlations within-modality were high and, to some extent, subject to FMC modulations as well. CBF and CMRO_2_ showed the strongest relationship overall (r = 0.96, all trials, q < 0.001), remaining FMC-independent across subgroups (r = 0.99 high FMC, r = 0.98 low FMC). SE-BOLD correlated moderately with GE-BOLD (r = 0.56, q < 0.001), CBF (r = 0.43, q < 0.001), and CMRO_2_ (r = 0.32, q < 0.01) in the full dataset. These relationships were fairly stable across FMC subgroups (Δz ≤ 0.18), though the SE-BOLD–CMRO_2_ correlation lost significance in the low FMC subgroup (r = 0.32, q > 0.05). The strongest FMC-dependent effect within the MRI parameters was observed for GE-BOLD: its correlation with CBF increased from 0.55 (all trials, q < 0.001) to 0.73 in the high FMC subgroup and dropped to 0.44 in the low FMC subgroup (Δz = 0.44), with a comparable pattern for GE-BOLD and CMRO_2_ (r = 0.34, q <0.01 to 0.67 high vs. 0.33 low; Δz = 0.48). Within the fNIRS metrics, HbO showed strong coupling with both HbT (r = 0.80, q < 0.001) and HbR (r = 0.72, q < 0.001) in the full dataset. Both relationships strengthened in the high FMC subgroup (r = 0.85 and 0.75) and weakened in the low FMC subgroup (r = 0.68 and 0.61; Δz = 0.42 and 0.27), whereas HbR–HbT remained stable across subgroups (r = 0.53, 0.52, 0.54; Δz = −0.03).

Results for the uncorrected-amplitude fNIRS data can be found in the Supplementary Material (Figure S4).

## 4. Discussion

This study provides, to our knowledge, the first quantitative comparison of CW-fNIRS chromophore modulations against the full suite of physiological parameters derived from GE–SE calibrated fMRI (GE-BOLD, SE-BOLD, CBF, CMRO_2_ and FMC) during a visual stimulation task. We characterised the relationship between optical and MRI signals, with particular attention to the interplay between haemodynamic and metabolic information and the vascular compartment each measurement reflects. Previous multimodal studies have incorporated fNIRS into fusion models to estimate CMRO_2_ without the calibrated-fMRI framework, relying instead on assumptions about baseline optical quantities (Torricelli et al., 2001) or other free model parameters (Huppert et al. 2008), or fused fNIRS with MRI models to estimate CMRO_2_ (Boas et al. 2003; Tak et al. 2010; Yucel et al. 2014), OEF (Hoge et al. 2005), or CBV (Alderliesten et al. 2014). Our approach here is different, as we aimed to characterise the cross-modal relationship itself, reporting the relationship between all measured physiological parameters and the link with the flow-metabolism coupling.

First, all three fMRI modalities detected robust, spatially consistent activation in the primary visual cortex, but with different spatial specificity and statistical sensitivity: GE-BOLD showed the largest clusters, SE-BOLD a more localised pattern, and CBF the smallest, in accordance with previous studies (Schmidt et al. 2005; Harmer et al. 2012). The magnitude of CBF and CMRO_2_ changes and the FMC values we observed during visual stimulation were also comparable to previous works (Hoge et al. 1999, Yücel et al. 2014). At both group and individual level, fNIRS chromophores followed the same temporal dynamics as the fMRI-derived signals, with HbO positively and HbR negatively correlated with fMRI signals across all modalities. This confirmed that the task-evoked response is reliably captured by optical measures at the population and single-subject level, supporting the use of fNIRS as a single-subject monitoring tool for clinical or naturalistic applications (Klein 2024).

The strong temporal correlations replicate and extend prior multimodal works (Huppert et al., 2006; Steinbrink et al., 2006; Strangman et al., 2002; Scarapicchia et al., 2017; Pereira et al., 2023) and show that this temporal agreement extends beyond BOLD to fMRI derived CBF and CMRO_2_ signals as well. For example, Huppert et al. (2006) reported relatively low correlations (< 0.3) between HbR and CBF for most of the subjects, whereas our correlation was 0.82, possibly because of a different experimental design with a longer task period.

In the trial-by-trial analysis, we sought to investigate whether the magnitude, not just the timing, of the response is shared across modalities. It revealed that correlations between fNIRS and fMRI metrics were not uniform but depended systematically on the FMC. While HbO relationship with the fMRI metrics showed little dependence on FMC, the correlations between HbR and fMRI metrics strengthened markedly when FMC was higher. This pattern was observed also for GE-BOLD with CBF and CMRO_2_ across the same subgroups, and this is reasonable as the GE-BOLD contrast arises directly from paramagnetic deoxyhaemoglobin (Ogawa et al., 1993). A first explanation for this pattern is a difference in SNR: HbR amplitudes are systematically smaller than HbO (Kinder et. al, 2022). In this perspective, trials with higher FMC would simply correspond to larger physiological variabilities in haemodynamic responses thus surpassing the noise level. We consider this explanation incomplete, however, since HbR mean (and median) modulations (Figure 4b) and HbO relationship with CBF and CMRO_2_ changes remained essentially stable across FMC subgroups. We therefore do not exclude the possibility that FMC indicates how tightly the deoxyhaemoglobin signal couples to flow, plausibly because large increases in flow relative to metabolism dilute local deoxyhaemoglobin concentration, linking HbR and GE-BOLD more tightly, while more balanced flow-metabolism changes weaken this link. The stronger correlations between fMRI signals and HbR with higher FMC can be interpreted in two non-mutually exclusive ways: on one side, HbR could be considered as a useful state-dependent proxy for metabolic or perfusion changes; on the other side, one could argue that HbO stably correlates with perfusion and could be better suited as a general proxy, given also the higher SNR. Additional multimodal work in conditions where BOLD changes oppose metabolism (Epp et al., 2025) could be beneficial to elucidate this. The relatively low cross-modal relationships for SE-BOLD across all FMC subgroups may reflect the intrinsically lower SNR relative to GE-BOLD. Alternatively, SE-BOLD might not outperform GE-BOLD as an fNIRS correlate because fNIRS signals themselves are sensitive to large vessels (Seddone et al., 2022).

FMC modulated similarly the GE-BOLD correlation with CBF and CMRO_2_ changes. Because CMRO_2_ modulation in this framework is itself derived algebraically from GE-BOLD and CBF changes via the Davis model, some portion of this modulation should be expected on mathematical grounds. The FMC-dependent relationship between GE-BOLD and CBF changes is also consistent with the Davis model: for a fixed CBF modulation, a higher BOLD signal implies a lower CMRO_2_ change in the model, thus a higher FMC. This has implications for any study that treats FMC as a fixed constant rather than a trial-varying quantity (Tak et al. 2010). Determining the FMC ratio is important for the quantitative interpretation of the BOLD signal itself: for a given change in CMRO_2_, different FMC values can produce different BOLD responses. Even if the specific magnitude of CMRO_2_ change depends on stimulus intensity and baseline conditions (Chiarelli et al. 2007), FMC itself has emerged as a characteristic of cerebrovascular physiology in the healthy human brain that could serve as a biomarker for certain neurological conditions (Tak et al. 2011).

### Limitations and Future Directions

First, CW-fNIRS cannot isolate baseline haemoglobin concentrations or separate absorption from scattering effects. As pointed out by previous studies (Uludağ et al., 2002, 2004; Strangman et al., 2003), this, combined with partial volume factors arising from the source detector positions relative to activated region and the subjects’ anatomical features can all potentially cause large variance in the fNIRS amplitude changes between subjects. We tried to mitigate this problem by correcting the fNIRS signal amplitudes based on their Jacobian, as computed from each subject’s own anatomy to partially counterbalance the assumption of a homogeneous head model.

The cross-modal comparisons described in this study focused on the temporal and quantitative aspect of the measurement but disregarded the spatial localisation of the changes. Cap placement was challenging and occasionally shifted downwards, introducing a slight misalignment of the measured ROI across subjects. Also, the fNIRS montage lacked the spatial density needed to produce a tomographic map comparable to the fMRI one. As a result, all the statistical analyses relied on metrics averaged on the occipital lobes for fMRI, and on selecting the most sensitive channels for fNIRS, thus sacrificing spatial granularity.

Additionally, the trial-by-trial subgroup analysis relied on a data-driven median split of FMC, a quantity itself derived from CBF and CMRO_2_ changes. As such, the specific threshold and grouping method used to define high- and low-FMC subgroups can influence the resulting correlation estimates, and trials near the group boundary or at the extremes of the CBF/CMRO_2_ distributions may disproportionately affect the observed subgroup differences.

Future studies should implement time-domain fNIRS alongside gas-free calibrated fMRI protocols to determine if absolute baseline optical quantifications improve cross-modal correlations, and whether they can be adopted in fusion models to obtain independent estimates of CMRO_2_ (Amendola et al. 2021). Also, different neural networks or patient populations with compromised neurovascular coupling (e.g., stroke or neurodegenerative diseases) should be investigated.

## 5. Conclusion

Both HbO and HbR showed strong and consistent temporal correlations with fMRI-derived haemodynamic and metabolic signals at both group and subject level. Trial-by-trial relationships between modulation amplitudes of the different modalities were significant but not constant, as they depended on the FMC. Both fNIRS chromophores and BOLD signals showed FMC-dependent correlations with CBF and CMRO_2_ changes, highlighting FMC as a key physiological variable that modulates the quantitative agreement between fMRI and fNIRS metrics. These findings offer rigorous grounding for fNIRS signal interpretation, demonstrating its utility as a surrogate marker for specific haemodynamic and metabolic parameters.

## Funding sources

This work was supported by the following funding bodies:

EU-NextGeneration EU- Italian Ministry of University and Research (MUR), National Plan for Recovery and Resilience (PNRR) Project Code: PE0000006, Project Title: “MNESYS-A multiscale integrated approach to the study of the nervous system in health and disease”. CUP: B83C22004960002.

GR was supported by EU-NextGeneration EU-MUR, Projects of National Relevance (PRIN). Project Code 2022MHMSSJ, Project Title: “Combining magnetic resonance imaging and optical spectroscopy to map microvascular function and oxygen metabolism in healthy and diseased brain”. CUP: D53C24004560006

The other authors were partially supported by:

1. EU-NextGeneration EU-MUR, PNRR, Mission 4 Component 2 –M4C2, Investment 1.5 – Call for tender No. 3277 of 30.12.2021 MUR Award Number: ECS00000041, Project Title: “VITALITY – Innovation, digitalization and sustainability for the diffused economy in Central Italy”, Concession Decree No. 1057 of 23.06.2022 adopted by MUR. CUP:D73C22000840006 .
2. EU-NextGeneration EU-MUR, Research National Program (PNR) and PRIN. Project Code: 2022BERM2F, Project Title: “Mapping Mitochondrial Function and Oxygen Metabolism in the Human Brain with Magnetic Resonance Imaging”. Funding call No.104 of 02.02.2022, Concession decree No. 1065 of 18.07.2023 adopted by MUR, ERC Panel LS7 “Prevention, Diagnosis and Treatment of Human Diseases”. CUP: D53D23013410001.
3. EU-NextGeneration EU- MUR, PNRR and PRIN, Project Code: P20225AEEE, Project Title: “Hybrid PET-MRI to simultaneously probe brain metabolism and cerebrovascular function in neurodegenerative diseases.” Funding call No. 1409 of 14.09.2022, Concession decree No. 1369 of 01.09.2023 adopted by MUR, ERC Panel LS7 “Prevention, Diagnosis and Treatment of Human Diseases”. CUP: D53D230 21480001.
4. European Union-NextGenerationEU (NGEU) – Italian Ministry of University and Research (MUR), National Plan for Recovery and Resilience (PNRR) and Projects of National Relevance (PRIN). Project number: P2022ESHT4, Project Title: “Advancing MRI biomarkers of brain tissue microstructure and energetics in Multiple Sclerosis.” Funding call No. 1409 of 14.09.2022, Concession decree No. 1367 of 01.09.2023 adopted by MUR, ERC Panel LS5 “Neuroscience and Disorders of the Nervous System”. CUP: D53D23019210001.

## Resource data for this article

The data that support the findings of this study are available from the corresponding author upon request. The code required to compute multiparametric maps using the model detailed in the article is available at https://github.com/chiarell/Hypercapnic-Calibrated-fMRI.

## Declaration of generative AI and AI-assisted technologies in the manuscript preparation process

During the preparation of this work the authors used Claude (Anthropic) in order to assist with reviewing texts and code. After using this tool, the authors reviewed and edited the content as needed and take full responsibility for the content of the published article.

