## Supplementary Material for "A comparison of continuous-wave fNIRS with quantitative fMRI-derived indices of brain function in the visual cortex"

**Figure S1**

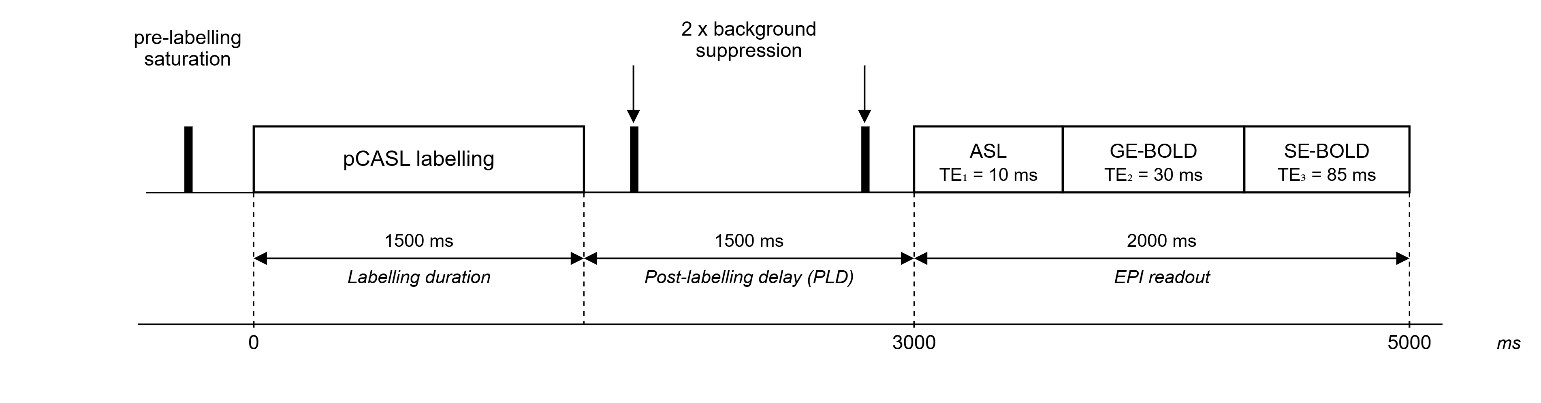

Figure S1: Pulse sequence timing diagram for gradient-echo spin-echo dual-excitation pseudo-continuous arterial spin labelling (GESE DEXI-pCASL) sequence.

**Table S1**

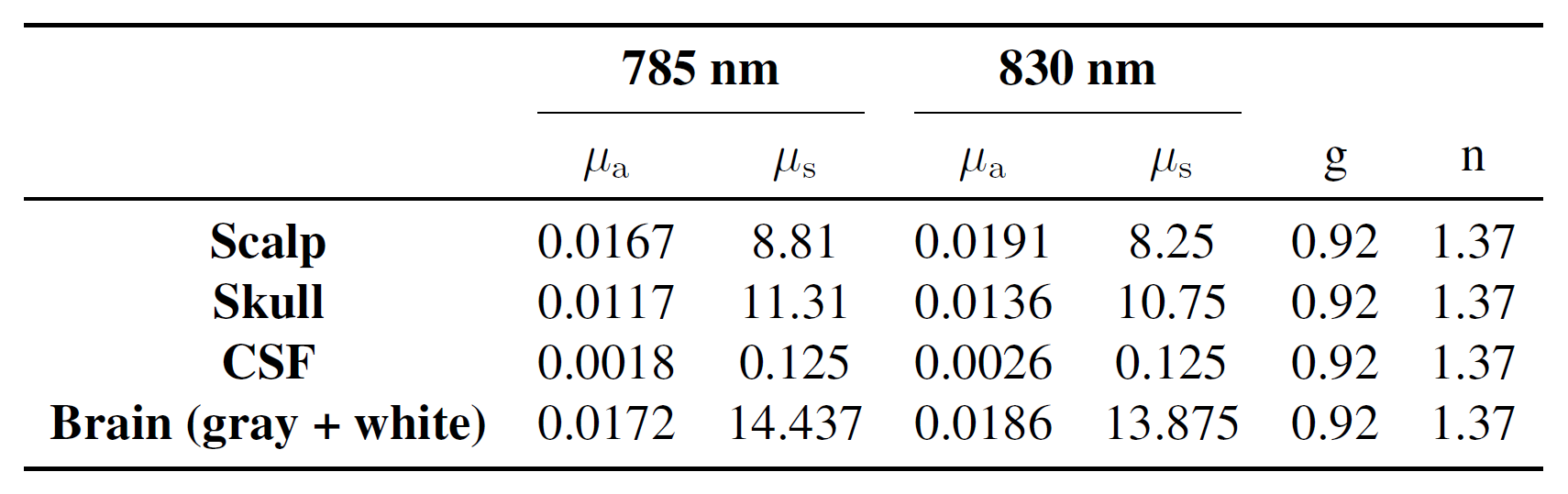

Table S1: Optical properties of the five-layered segmented head tissue for forward light problem with Monte Carlo simulations (Strangman et al., 2003): absorption coefficient μ_a_(mm^−1^), scattering coefficient μ_s_(mm^−1^), anisotropic factor g and index of refraction n.

**Table S2**

| Sub # | % Good channels |
| --- | --- |
| 01 | 100% |
| 02 | 84% |
| 03 | 42% |
| 04 | 97% |
| 05 | 98% |
| 06 | 81% |
| 07 | 69% |
| 08 | 81% |
| 09 | 24% |
| 10 | 32% |
| 11 | 66% |
| 12 | 95% |
| 13 | 95% |

Table S2: fNIRS data quality results for each channel. The % of good channels was calculated after discarding the channels with source-separation greater than 45 mm. Subjects with less than 60 % good channels were discarded.

**Table S3**

|  | **H** | **k** | **x** | **y** | **z** | **t** |
| --- | --- | --- | --- | --- | --- | --- |
| **GE BOLD** |  |  |  |  |  |  |
| Right inferior occipital gyrus | R | 155 | 48 | -72 | -16 | 20.14 |
|  |  |  | 34 | -74 | -22 | 8.52 |
|  |  |  | 32 | -78 | -14 | 6.94 |
| Left inferior occipital gyrus | L | 484 | -24 | -96 | -2 | 16.79 |
|  |  |  | -12 | -96 | 14 | 14.79 |
|  |  |  | -10 | -102 | 6 | 8.04 |
| Right lingual gyrus | R | 41 | 8 | -88 | 14 | 8.84 |
| Left inferior occipital gyrus | L | 574 | 18 | -98 | -4 | 8.73 |
|  |  |  | 22 | -100 | -12 | 8.11 |
|  |  |  | 30 | -92 | 0 | 8.08 |
| Left occipital fusiform gyrus | L | 32 | -28 | -68 | -14 | 7.38 |
| Right superior occipital gyrus | R | 48 | 28 | -76 | 14 | 7.04 |
| **SE BOLD** |  |  |  |  |  |  |
| Left inferior occipital gyrus | L | 420 | -28 | -96 | 0 | 15.26 |
|  |  |  | -20 | -102 | -4 | 10.72 |
|  |  |  | -8 | -98 | -12 | 9.72 |
| Right occipital pole | R | 580 | 28 | -90 | -6 | 14.64 |
|  |  |  | 14 | -102 | 10 | 11.16 |
|  |  |  | 34 | -88 | 8 | 10.42 |
| **CBF** |  |  |  |  |  |  |
| Right occipital pole | R | 107 | 16 | -102 | -4 | 7.15 |
|  |  |  | 36 | -88 | 12 | 6.38 |
|  |  |  | 30 | -96 | -2 | 6.31 |
| Left inferior occipital gyrus | L | 10 | -24 | -94 | 4 | 5.98 |
| Left occipital pole | L | 15 | -22 | -102 | -10 | 5.30 |
|  |  |  | -24 | -104 | -2 | 4.67 |
| Left inferior occipital gyrus | L | 9 | -32 | -98 | -2 | 5.16 |

Table S2: Significantly activated regions at the group level for GE-BOLD, SE-BOLD and CBF. The statistical threshold was defined as p < 0.05 FWE corrected for multiple comparisons at cluster level with an extent threshold of 10 voxels for GE-BOLD and SE-BOLD, and p < 0.001 with an extent threshold of 5 voxels for CBF. For each cluster, the region with the maximum t-value is listed first, other regions in the cluster are listed below. The hemisphere (H) and the number of voxels (k) are shown for each cluster. The MNI coordinates (x,y,z) in mm and the t-value (t) are provided for each peak voxel.

**Section S1: Data-driven simulations for fNIRS channel selection**

In order to select the fNIRS channels to be compared with the corresponding fMRI data for each subject, we considered the Jacobian of each channel within the activated fMRI region. After summing the Jacobian across active vertices, we obtained a single Jacobian value for each channel. Two strategies were evaluated: retaining only channels with a Jacobian value greater than a certain threshold (in our case, 50% of the maximum Jacobian value for each subject) or selecting the *n* (2 to 6) channels with the highest Jacobian values. The first strategy ensured high sensitivity in the activated region, but resulted in a different number of channels selected for each subject. The second strategy ensured the same number of channels across subjects, despite potentially lower sensitivity. We explored these options: two representative subjects are shown in Figure S2a. The averaged group-level time-series (Figure S2b) did not differ substantially in shape, showing essentially lower amplitude as the number of selected channels increased, since the likelihood of introducing noisy channels was higher. We chose to consider the top two channels for further analysis, since the difference in the number of selected channels across subjects for Jacobian > 50% max was marked (e.g. 1 channel vs 5).

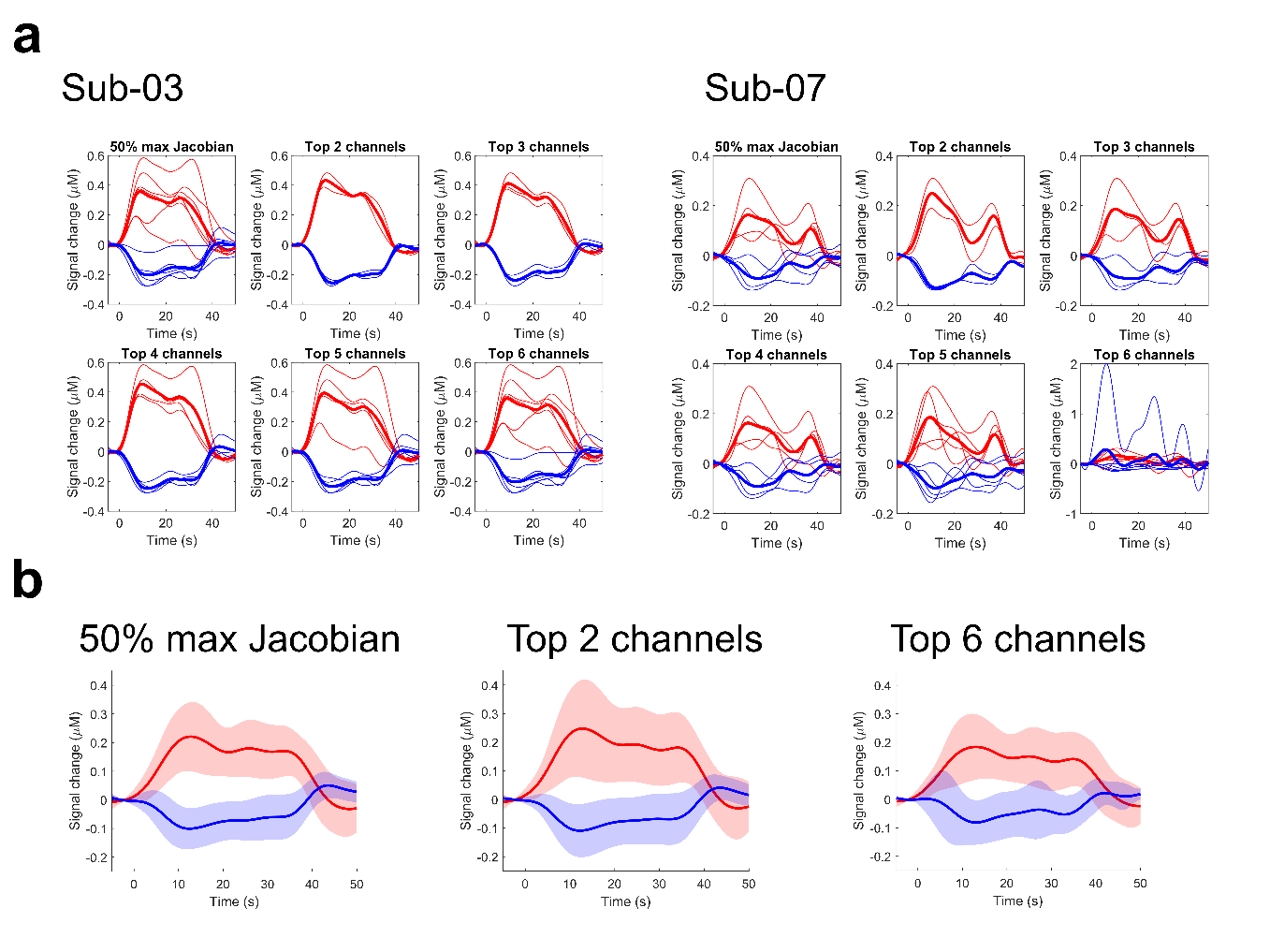

Figure S2: **Channels selection strategies for fMRI coregistration**. (a) All strategies are shown for two representative subjects, respectively channels with Jacobian values higher than 50% of the maximum Jacobian, and the 2 to 6 channels with the highest Jacobian. The average of the selected channels is shown with a bold line for each plot. (b) Averaged group-level time-series different strategies. Solid lines represent the mean across subjects; shaded areas indicate ± standard deviation.

**Section S2: fNIRS sensitivity-based amplitude correction**

MP2RAGE images were segmented using the SIAM model, selected for its accuracy in scalp and skull delineation. The SIAM segmentation provides 17 anatomical labels. Since Monte Carlo light propagation simulations were performed using a simplified 5-tissue model (scalp, skull, CSF, gray matter, white matter; Figure S3a), several labels were merged for consistency. Cerebellar gray matter, thalamus, pallidum, putamen, caudate, nucleus accumbens, amygdala, and hippocampus were merged into the gray matter label; ventricular CSF, vessels, and dura mater were merged into the CSF label. Scalp and cortical surface meshes were extracted using CAT12 segmentation, while the skull surface mesh was generated from the SIAM segmentation. For each source-detector channel, the volumetric sensitivity (Jacobian) computed from Monte Carlo simulations was interpolated onto three surface meshes: scalp, skull, and cortex (Figure S3b). This yielded a Jacobian value J(v) at each vertex v of the three meshes. Vertices with Jacobian values below 1% of the maximum value across all three surfaces were discarded. The resulting spatial distribution shows the expected pattern of decreasing sensitivity with depth, with the highest values on the scalp and the lowest on the cortex (Figure S3c). For each channel, the total cortical sensitivity was computed as the sum of Jacobian values over all vertices belonging to the cortical surface:

$$J_{\text{cortex}}=\sum_{v\in\mathcal{V}_{\text{cortex}}} J\left( v \right)$$

and the total sensitivity across all three tissue surfaces was computed as:

$$J_{\text{total}}=\sum_{v\in\mathcal{V}_{\text{cortex}}\cup\mathcal{V}_{\text{skin }}\cup\mathcal{V}_{\text{skull}}} J\left( v \right)$$

The correction factor for the channel was then defined as the ratio of cortical to total sensitivity:

$$c=\frac{J_{\text{cortex}}}{J_{\text{total}}}$$

Across all included channels, c ranged approximately between 0.1 and 0.4, reflecting the proportion of total light sensitivity attributable to the cortical layer relative to the extracerebral tissues. Each channels amplitude S(t) was then divided by c:

$$S_{\text{corr}}\left( t \right)=\frac{S\left( t \right)}{c}$$

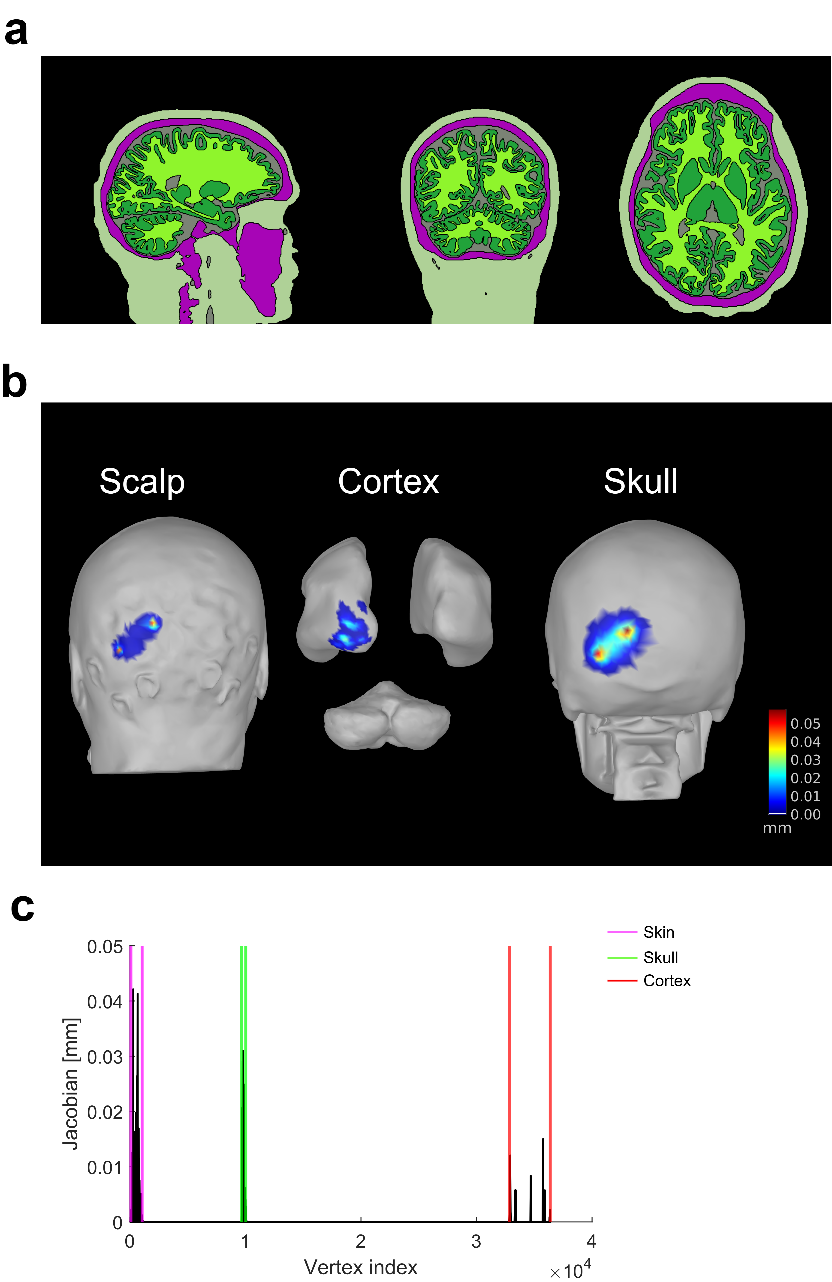

Figure S3: **Sensitivity-based correction pipeline for fNIRS signal amplitude using multi-layer Jacobian interpolation.** (a) Sagittal, coronal, and axial views of a representative segmented MP2RAGE image with the SIAM model, showing the tissue layers used for volume-to-surface interpolation: scalp (light green), skull (purple), CSF (grey), grey matter (dark green), and white matter (bright green). (b) Representative single-channel Jacobian sensitivity maps projected onto the scalp, cortex, and skull surfaces. (c) Jacobian sensitivity values plotted for all mesh vertices for the same channel, with vertical lines marking the boundaries between the skin (magenta), skull (green), and cortex (red) mesh segments.

**Figure S4**

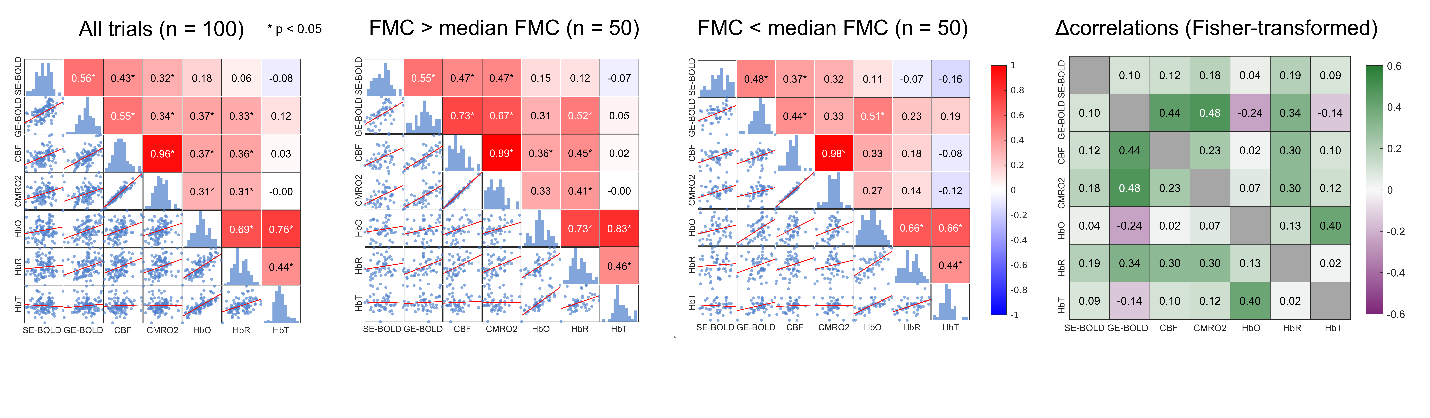

Figure S4: **Trial-by-trial response amplitudes and cross-modal quantitative correlations for uncorrected (amplitude) fNIRS data.** Pairwise Pearson correlation matrices of single-trial beta weights for all signal pairs, computed across all trials (left), high FMC trials (FMC > median, n = 50, center), and low FMC trials (FMC < median, n = 50, right). Asterisks indicate statistically significant correlations (p < 0.05). Scatter plots below the diagonal show individual trial data; histograms on the diagonal show each signal's marginal distribution. The matrix on the right represents the difference between the Fisher-transformed correlations of the high and low FMC groups.
